# Ultra-fast protein structure prediction to capture effects of sequence variation in mutation movies

**DOI:** 10.1101/2022.11.14.516473

**Authors:** Konstantin Weissenow, Michael Heinzinger, Martin Steinegger, Burkhard Rost

## Abstract

Top protein three-dimensional (3D) structure predictions require evolutionary information from multiple-sequence alignments (MSAs) and deep, convolutional neural networks and appear insensitive to small sequence changes. Here, we describe *EMBER3D* using embeddings from the pre-trained protein language model (pLM) ProtT5 to predict 3D structure directly from single sequences. Orders of magnitude faster than others, EMBER3D predicts average-length structures in milliseconds on consumer-grade machines. Although not nearly as accurate as *AlphaFold2*, the speed of EMBER3D allows a glimpse at future applications such as the almost real-time rendering of deep mutational scanning (DMS) movies that visualize the effect of all point mutants on predicted structures. This also enables live-editing of sequence/structure pairs. EMBER3D is accurate enough for highly sensitive rapid remote homology detection by *Foldseek* identifying structural similarities. Overall, our use cases suggest that speed can complement accuracy, in particular when accessible through consumer-grade machines. EMBER3D is free and publicly available: https://github.com/kWeissenow/EMBER3D.

## Introduction

### AlphaFold2 advances protein structure prediction

The Critical Assessment of protein Structure Prediction (CASP) has provided the gold-standard to evaluate protein structure prediction for almost three decades ^1^. At its first meeting (CASP1 Dec. 1994), the combination of machine learning (ML) and evolutionary information derived from multiple sequence alignments (MSAs) reported a major breakthrough in secondary structure prediction ^2^. 2021’s method of the year ^3^, *AlphaFold2* ^4^ has combined more advanced machine learning (ML) with larger MSAs and more potent hardware to substantially advance protein structure prediction. But even this pinnacle of 50 years of research has a shortcoming: predictions are mostly insensitive to small variations in the input ^5^.

### Protein language models (pLMs) substitute evolutionary information

Advances in natural language processing (NLP) spawned protein language models (pLMs) ^6-11^ that leverage the wealth of information in exponentially growing but unlabelled protein sequence databases by solely relying on sequential patterns found in the input. Processing the information learned by such pLMs, e.g., by inputting a protein sequence into the network and constructing vectors from the activation in the network’s last layers, yields a representation of protein sequences referred to as embeddings (Fig. S1 ^11^). This allows to transfer features learned by the pLM to any downstream (prediction) task requiring numerical protein representations (transfer learning) which has already been showcased for various aspects of protein prediction ranging from structure ^5,12^ and disorder ^13^ to function ^14^. Distance in embedding space correlates more with protein function than with sequence similarity ^15^ and can help clustering proteins into families ^16-18^. Recently, pLMs have been used to directly predict protein 3D structure ^5,19-22^. Using embeddings from pLMs instead of evolutionary information from MSAs simplifies and speeds up structure prediction. Speed is gained at the price of accuracy while precomputed *AlphaFold2* predictions are available for over 200 million proteins ^23^. Is there a value in gaining speed even when losing accuracy?

Here, we introduce a novel solution using sequence embeddings and attention heads (AHs) from ProtT5 ^11^ to predict inter-residue distances (2D structure) and backbone atom coordinates (3D structure). Without any MSA, our proposed solution reaches competitive performance at unprecedented speed. Paired with its sensitivity to small changes in the input, we showed that the proposed solution opens the door for novel applications such as the generation of “*protein mutation movies*” (PMMs). Each frame in these movies shows the predicted structure of a mutant. Connecting these frames, the rendered movie visualizes the mutational landscape of a protein. We also showed that our prediction quality sufficed for fast structure alignment, albeit not reaching the performance of AlphaFold2-like solutions.

## Results

### Ultra-fast: protein structure predicted in sub-seconds

We benchmarked the prediction speed of EMBER3D through poly-alanine sequences with 20-1500 residues (i.e., alanines). EMBER3D predicted backbone atom coordinates (3D structure) and inter-residue distances (2D structure) in sub-seconds for average length sequences (Fig. 1B, e.g., 0.3s for 384 residues). In contrast, ESMFold ^21^ needs 14.2s (factor 47) for the same sequence, i.e. EMBER3D was as much faster than ESMFold as that sped up over AlphaFold2. Additionally, our model is small enough to fit onto consumer-grade GPU hardware (e.g., NVIDIA Geforce 1080ti with 8 GB of video memory) which can predict for most proteins (<430 residues). On server-grade hardware (e.g., NVIDIA Quadro RTX 8000 with 48 GB), proteins of up to ^∼^700 residues are predicted in less than a second while the maximal length rises to 1420 residues (Fig. 1B). To put this into perspective, we applied the publicly available implementation *ColabFold* ^24^ speeding up *AlphaFold2* ^4^ and two newer 3D prediction methods using pLMs, namely *OmegaFold* ^20^ and *HelixFold* ^19^ on our test set (CASP14 domains ^25^) using the same hardware (NVIDIA Quadro RTX 8000). EMBER3D significantly outpaced the others (Fig. 1A). Its speed allowed predicting structures for 538,488 proteins from the entire Swiss-Prot database ^26^ in just 60.5 hours on a single server-grade GPU. Since structural prediction is obtained near-instantaneously our solution allows live-editing of sequences.

**Fig. 1.**
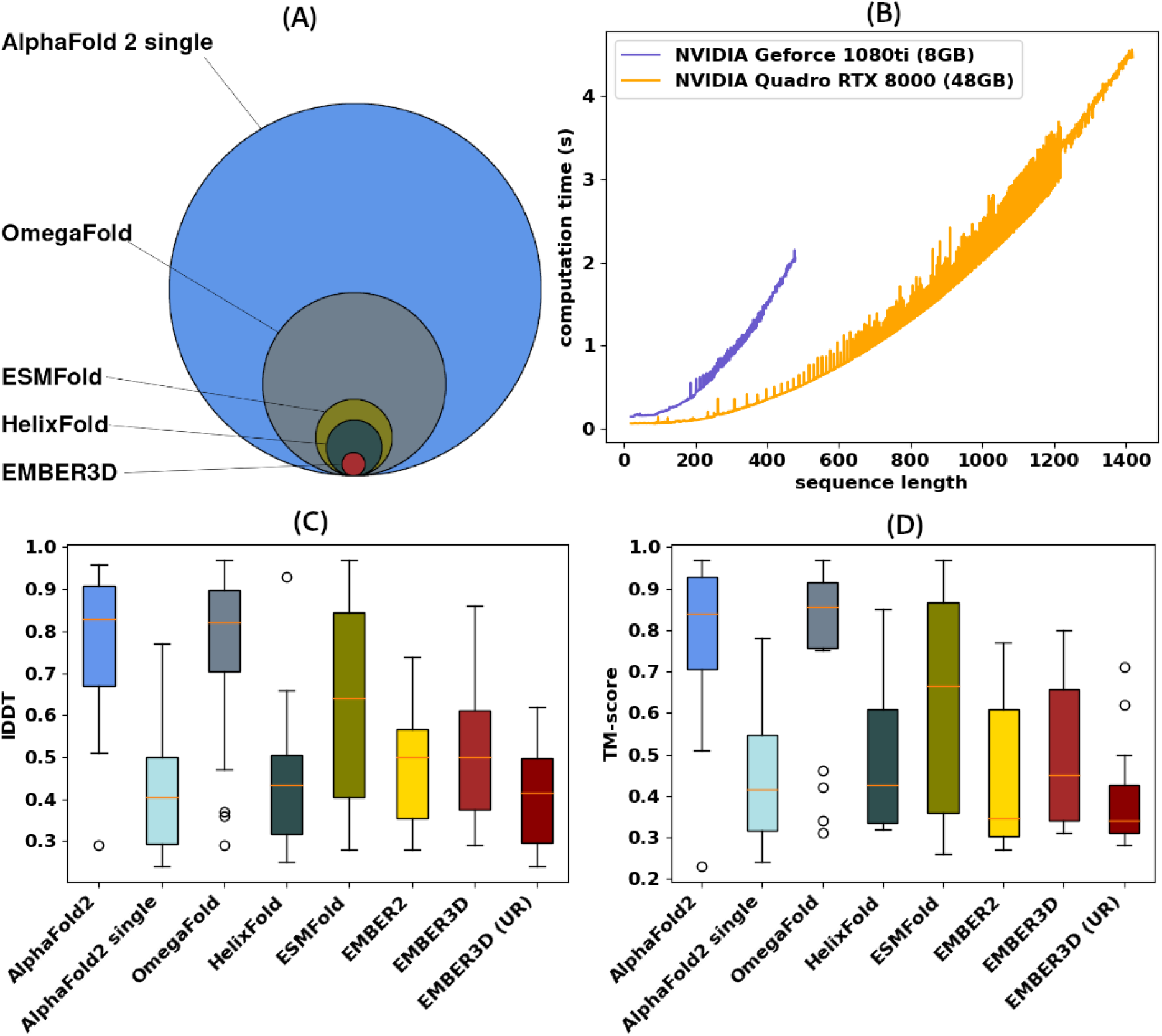
Protein structure predicted in milliseconds. **(A)** Relative comparison of inference runtime of *AlphaFold2* ^4^ (inputting single sequence instead of multiple sequence alignments - MSAs), *OmegaFold* ^20^, *ESMFold* ^21^, *HelixFold* ^19^, and *EMBER3D* on the test set of *CASP14* domains^52^. Circle areas are proportional to the respective fraction of the runtime of AlphaFold2 (actually proxied by its faster derivative *ColabFold* ^24^). **(B)** Most proteins can be predicted by EMBER3D on consumer-grade hardware although runtime (y-axis in seconds) grows exponentially with protein length (y-axis). EMBER3D predicts structures of proteins <700 residues in less than a second, orders of magnitude faster than competing methods. Its model is small enough to allow predictions for average-length proteins even on consumer hardware (purple). Prediction accuracy measured with local Distance Difference Test (IDDT)^27^ **(C)** and TM-score^58^ (**D**) on our test set (CASP14 domains^52^) for *AlphaFold2* (AlphaFold2 sped- up through ColabFold^24^), *AlphaFold2 single* not using MSAs as input but only the query sequence, *OmegaFold, HelixFold, ESMFold, EMBER2* ^5^ and the method introduced here *EMBER3D* (relaxed with pyRosetta and unrelaxed (UR) backbone output). While AlphaFold2 clearly outperformed the EMBER family, prediction quality dropped significantly when disregarding evolutionary information by only using single sequence input.

### Single-sequence predictions better than for AlphaFold2

In order to assess the quality of EMBER3D predictions, we compared TM-score and lDDT^27^ on our test set (CASP14 domains) with results from ESMFold, OmegaFold, HelixFold, our previous method, EMBER2, and AlphaFold2 with MSAs (standard) and without using MSAs as input (Fig. 1CD). Although AlphaFold2/ColabFold significantly outperformed the EMBER family, performance dropped substantially without MSA input (Fig. 1CD: “AlphaFold2 single”). While HelixFold performed similar to the EMBER family, OmegaFold outperformed all sequence-based methods, including ESMFold, albeit at significantly higher runtime (Fig. 1A).

### EMBER family more sensitive to mutants (SAVs) than AlphaFold2

In contrast to methods relying on MSAs, we hypothesized our single sequence-based method to be more sensitive to small sequence variations, e.g., to single amino acid variants (*SAVs*). We measured the structural effect of SAVs by computing the changes in structures predicted between wild type and mutated sequence measured in lDDT. Since EMBER3D outputs 2D and 3D structure, we considered both. We correlated predicted structural changes with deep mutational scanning (DMS) data for nine proteins ^28-34^ shorter than 250 residues. For AlphaFold2, our restricted resources forced to consider only the shortest five (loading our hardware for over a week). The SAV analysis required predicting 3D structures for all possible point mutants in a protein (e.g. 6,650 predictions for a protein of 350 residues). EMBER3D, completed this job within minutes (almost 10,000 times faster than ColabFold which is faster than AlphaFold2).

The differences between native and mutant structure predicted by AlphaFold2 correlated only weakly with DMS data (correlation over 5 proteins: 0.15, Fig. 2), significantly less than differences predicted by EMBER3D (Fig. 2). Amongst the EMBER3D methods, differences in predicted distance maps correlated more (0.36) than the 3D output (0.24, Fig. 2). While pLM-based ESMFold correlated better with DMS than AlphaFold2 (0.18), it did not reach EMBER3D on any sample, and OmegaFold remained even below (correlation over 5: 0.05). Only HelixFold outperformed the EMBER-family for one sample (Ubiquitin), despite a low overall correlation (0.18). Family-averaged AlphaFold2 and the protein-specific *EMBER*-family differed on average and in detail, e.g., the DMS set predicted by far best by AlphaFold2 (*SUMO-conjugating enzyme UBC9*) was below average for EMBER3D.

**Fig. 2.**
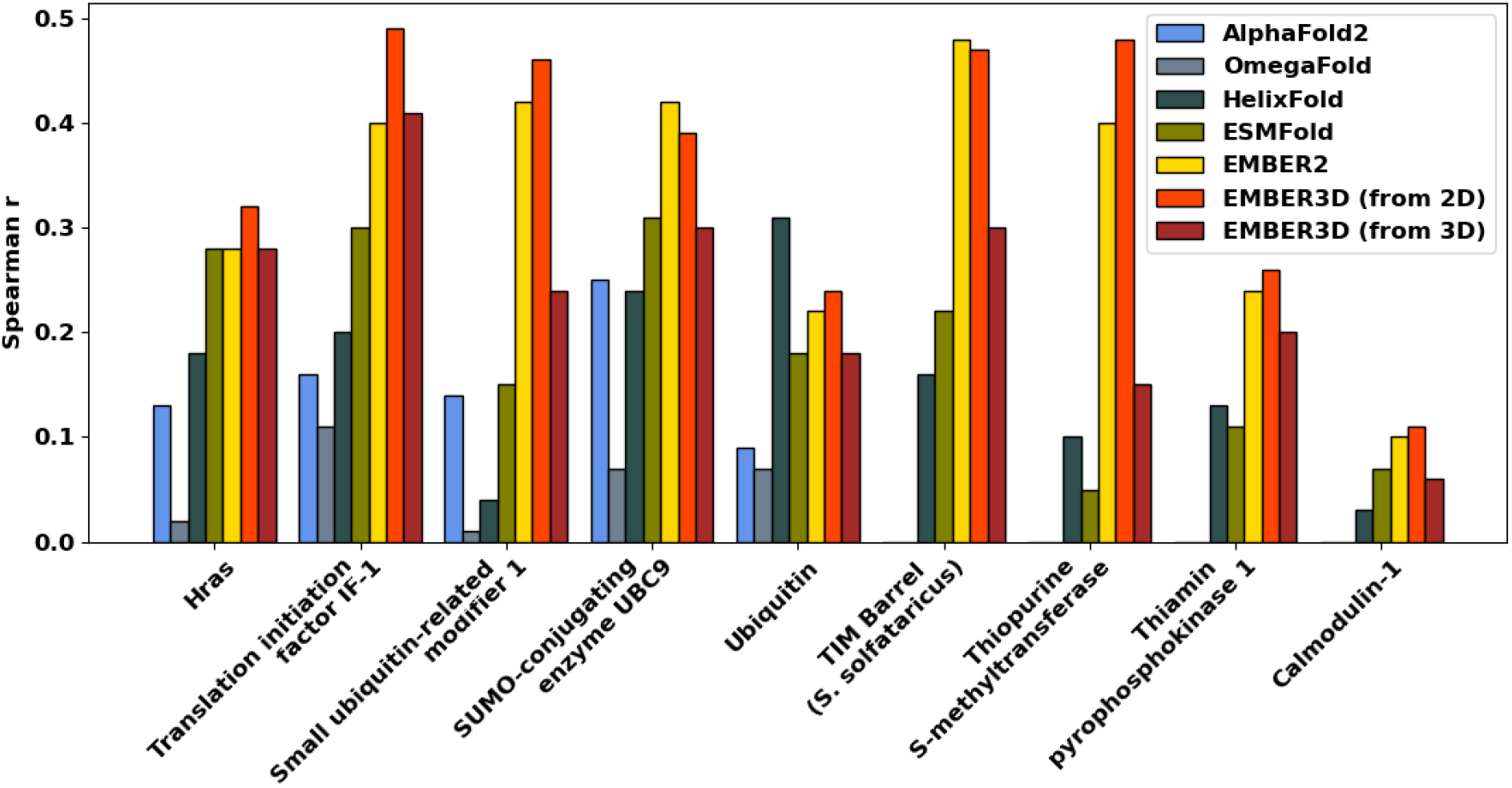
EMBER3D sensitive to point mutants (single amino acid variants, SAVs). y-axis: Spearman rank correlation between structural differences (measured in lDDT^27^) of wild type sequence and all possible amino-acid substitutions for nine proteins from a deep mutational scanning dataset (DMS^28-34^) for AlphaFold2^4^ (blue; only shortest five proteins due to constrained resources), OmegaFold^20^ (gray; only shortest five), HelixFold^19^ (dark green), ESMFold^21^ (green), EMBER2^5^ (yellow) and EMBER3D (orange, red). For EMBER3D we measured correlations using the lDDT computed from both the 2D (distance maps) and 3D (C-beta co-ordinates) outputs of the model, showing that the former more reliably captures DMS experiments. Overall AlphaFold2 and OmegaFold predictions correlated at best weakly with DMS data while EMBER3D (in particular with its 2D mode) correlated most.

### Protein mutation movies capture differences in predicted structures

To visualize the mutational landscape of a protein, we developed a tool which renders the differences between structures predicted for wild type and all possible point mutants (SAVs: length*19) into a protein *mutation movie (PMM)*. This tool first runs EMBER3D to obtain both 2D and 3D structure predictions for all possible SAVs in a protein sequence. Next, the tool generates three images for each mutant (SAV): (1) distance map for SAV (using matplotlib ^35^; Fig. 3C), (2) sketch of 3D backbone coordinates for SAV from a static viewpoint (using PyMol ^36^; Fig. 3B), and (3) a mutation profile (Fig. 3D). The tool then composes these three into one image (for each mutant), and renders the individual images as an animated movie clip using *ffmpeg* ^37^ with 19 frames per second, thereby showing all 19 possible amino-acid substitutions for one residue position in each second of playback time (Fig. 3).

**Fig. 3.**
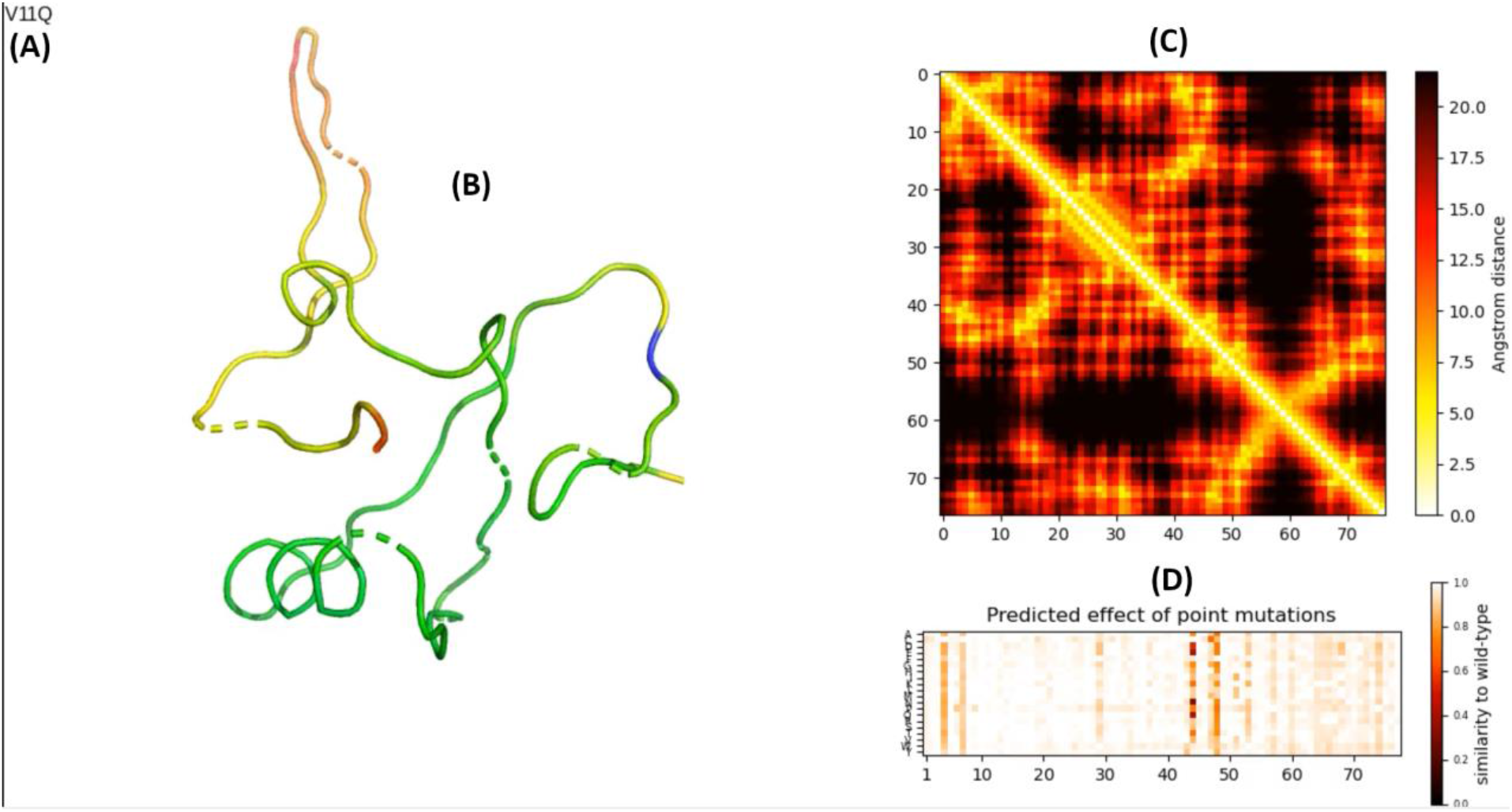
Screenshot of one from a protein mutation movie (PMM). **(A)** The top-left corner indicates the currently shown mutation (here V11Q implying that the valine V at position 11 is replaced by a glutamine Q). All structure predictions (panels B and C) are obtained for this single amino acid variant (SAV). **(B)** Shows the predicted 3D backbone structure of the mutant as a simplified sketch in which the coloring indicates the predicted confidence of the model (highest quality: green, lowest: red; blue marks the currently mutated position, here the 11-th residue). **(C)** Shows the predicted 2D structure (distance map) for the same mutant (V11Q; this map is trivially symmetric (distance(i,j)=distance(j,i)). **(D)** Presents the mutation profile for the entire protein, in analogy to the concept of DMS^28^ this view highlights the effect of all 19 non-native SAVs on the predicted structure (with respect to wild type) for all residues. Darker colors indicate a larger deviation of the mutant’s predicted structure from the wild-type.

### Predicted confidence correlates between EMBER3D and AlphaFold2

Copying ideas introduced by AlphaFold2 ^4^, EMBER3D also predicted its own reliability. Despite reaching lower overall performance than AlphaFold2, EMBER3D succeeded in assessing its own success (Fig. S2A). Although different in methodology and focus, we expected AlphaFold2 and *EMBER3D* to share the same trend in their ability to accurately predict structures for difficult proteins. Indeed, the predicted lDDT (pLDDT) confidence scores for 538,488 proteins from Swiss-Prot Spearman rank correlated to 0.53 (p-value ^∼^0.0).

EMBER3D, optimized for speed, was overall less confident than AlphaFold2 in its own predictions. However, most proteins for which AlphaFold2 predicted a high-quality structure were also predicted with higher-than-average confidence by EMBER3D (Fig. S2B). This suggested using EMBER3D as a rough, but very fast pre-for AlphaFold2.

### Foldseek paired with EMBER3D beats MMSeqs2

Protein structure predictions can unravel relations between proteins with highly diverged sequences^38^. We benchmarked the detection of *SCOP* folds from EMBER3D predictions by predicting structures for 11,211 domains from *SCOP-40* (v2.01) ^39^ and aligning these all against all with *Foldseek* ^40^. Predicted structures constituted the queries, experimental structures the inference lookup set ^40,41^. Replicating the *Foldseek* benchmark ^40^, we evaluated performance by measuring the sensitivity up to the 5th false positive per query for the three different levels of *family, superfamily*, and *fold*. Protein pairs with experimental structures were detected best by Foldseek (Fig. 4: sensitivity: 0.901, 0.578, 0.154 for family, superfamily and fold, respectively) followed by comparing EMBER3D-predicted pairs of structures (Fig. 4: 0.865, 0.499, 0.100), while sequence-sequence comparisons with *MMseqs2* were substantially less successful (Fig. 4: 0.543, 0.082, 0.002).

**Fig. 4.**
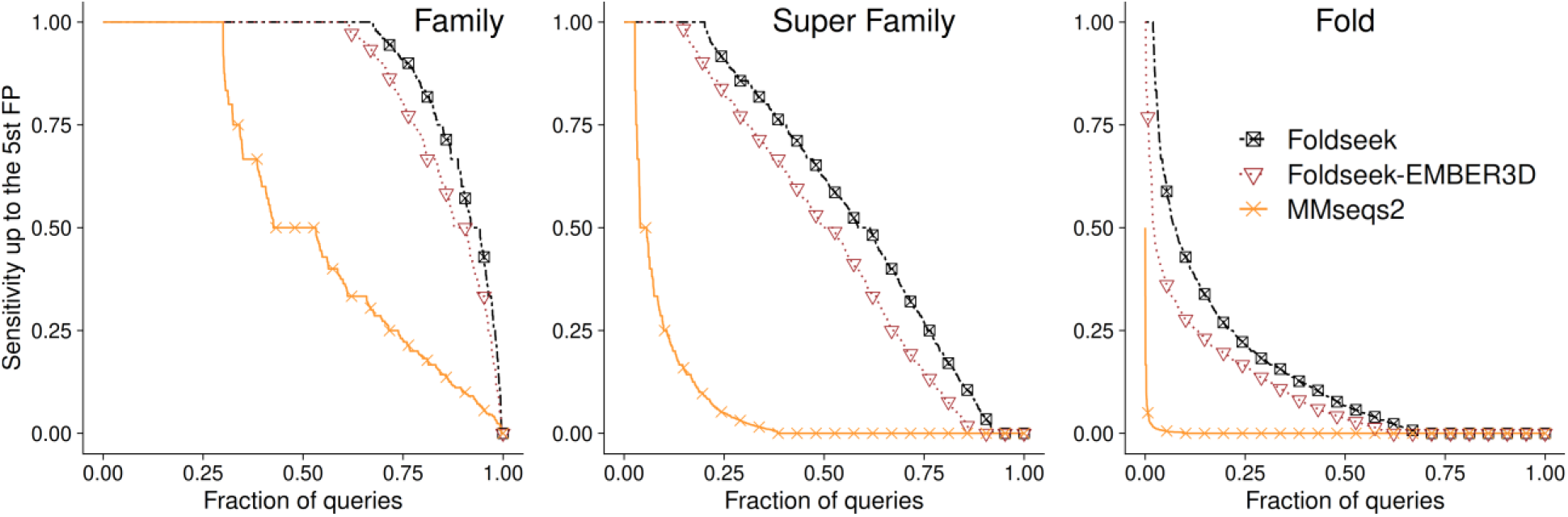
Foldseek on EMBER3D predictions significantly outperformed MMSeqs2. One way to profit from 3D predictions is to search with the structure of a query Q for other proteins with similar structure and potentially different sequences. *Foldseek*^*40*^ addressed this theme by making a fast structure-structure comparison available that clearly outperforms sequence comparisons when assessed on the three commonly used levels of increasing evolutionary distance, namely *Family* (most closely related), *Super Family* (more distantly related), and *Fold* (diverged sequences with similar structures). Here, we showed the *sensitivity up to the 5th false positive* (FP, y-axes) versus the *fraction of queries* (x-axes), which was the fraction in the subset of a dataset with 11,211 SCOP-40 domains^39^ (the actual numbers differed between the three panels/categories). Foldseek comparisons using EMBER3D predictions (red triangles) significantly outperformed alignment in sequence space with MMSeqs2^41^ (orange crosses) and almost reached the sensitivity of Foldseek applied to high-resolution experimental structures (black squares).

## Discussion

### Speed enables mutation movies for higher order mutants

*AlphaFold2* ^4^ and *RoseTTAFold* ^42^ continue to mark the state-of-the-art in protein structure prediction by using evolutionary information derived from multiple sequence alignments (MSAs). *ColabFold* ^24^ speeds up the sequence search without losing performance. *HelixFold* ^*19*^, *OmegaFold* ^*20*^, *ESMFold* ^*21*^, and RGN2 ^22^ speed up by replacing evolutionary information with embeddings from protein Language Models (pLMs). *EMBER3D* pushes much further: where ESMFold speeds up 10-60 times, EMBER3D adds more than another order of magnitude, even on less advanced hardware. Still, through its speed-up, ESMFold tripled the number of predictions to cover 600 million proteins ^23,43^. Instead of *going big*, we explored alternative ways enabled by our immense additional speed-up. For instance, we introduced the concept of “protein mutation movies” (PMMs, Fig. 3) allowing interactively to explore the predicted effects of point mutations (single amino acid variants: SAVs) upon 3D structure (Fig. 2).

The PMMs can benefit from EMBER3D’s speed only because, although less accurate than AlphaFold2, it better predicts effects of sequence variation (Fig. 2). Possibly because AlphaFold2, RoseTTAFold and similar solutions, that began with the PHD first successfully combining machine learning and MSAs ^2^, learn family-averages rather than protein-specific predictions potentially rendering results less sensitive to small changes in the input (Fig. 2). Also using pLM embeddings, ESMFold clearly surpasses EMBER3D’s accuracy for native sequences. However, ESMFold performed significantly worse in predicting the effect of point mutations (Fig. 2). This might result from the strong regularization of EMBER3D, i.e., its relatively tiny model with 4.7M trainable parameters (Table S1). As a result, small changes in the input are maintained throughout the network without getting mapped to the most similar structural motif seen during training. Even if ESMFold were more sensitive to SAV effects (Fig. 2), it still would not be fast enough to explore a large space of mutants (Fig. 1A), especially when considering to explore higher-order effects such as SAV duplets, triplets, asf.

As proof-of-principle, we pursued several examples of proteins by iteratively picking the SAV that changed the predicted structure most. This naïve heuristic already showed promising results but exploring different sampling strategies to navigate the energy landscape of mutants will be an important future research direction. In this context, the question will not really be how much speed-up over AlphaFold2/ColabFold suffices for *PMMs*, but which aspect becomes crucial first: either the lack of resources to optimize another N-th iteration (with N=3 for triplets) or the limited accuracy in predicting structure (errors might somehow cancel out for N>1 or might increase by error to the Nth power).

### Speed may not trump accuracy

ESMFold speeds up over AlpaFold2 at some costs to performance, EMBER3D speeds up more at more costs; random predictions would be instantaneous at higher loss. At which loss of performance is which gain in speed still a trump? There is no proxy for such a point, instead the answer depends on what the method is used for. PMMs are just one aspect. Another aspect is the usage of predicted structure to unravel unsuspected relations between protein pairs through the help of methods such as Foldseek ^40^ (Fig. 4). In order to leverage the improved sensitive of structure-over sequence-based search methods on a large scale, speedy structure predictions will be crucial as one needs to keep pace with the increasingly growing sequence databases. Another consideration could be the costs to the environment, e.g., by calculating the costs in terms of energy consumption penciling in the hardware and hardware-maintenance costs. Unfortunately, this is not well-defined because the speed advantage of pLM-based solutions shrinks non-linearly with protein length. Consequently, the gain will be much higher for predicting five bacterial than for the human proteome although both may have similar numbers of proteins, and they will be much higher for kinases than for structural proteins.

### What makes EMBER3D faster?

Presumably, the most obvious factor for EMBER3D outpacing others was the model with relatively few free parameters (Table S3). However, the total parameter count might suggest our model to be relatively slow, given its size (1.5B parameters). However, the underlying pLM, ProtT5 ^11^, accounts for 99.7% of those parameters, leaving only 0.3% (4.7M) for the actual structure prediction module. The frozen pLM (no gradients are backpropagated to the pLM, i.e., it is only used as static feature encoder), however, makes up only a small fraction of the compute time, especially, as it is run in half-precision, turning the structure module into the bottleneck for speed. Therefore, we kept this part of the model as small as possible. We further benefited from an optimized version of the SE(3) module provided by Nvidia that allowed to roughly half inference time at reduced memory consumption. We also reduced the computational complexity of the structure prediction module by only modeling backbone atoms. While neglecting side-chains loses information, for some use-cases the resulting speed-up compensated for this loss.

### Better for proteins from small families?

Although *EMBER3D* is much less accurate than AlphaFold2 (Fig. 1) and ESMFold ^21^ and RGN2 ^22^ are a little less accurate ^21^, all three appear to outperform MSA-based methods when tested on single sequence (Fig. 1). However, strictly speaking, at least for ESMFold and EMBER3D, this claim was a proxy: we simply tested what would happen if we took the proteins for which we do have large MSAs and pretended that we do not have those. We had to refrain to this proxy because there are no high resolution structures available for proteins without MSAs because experimental structural biologists successfully optimize the leverage of each structure ^44^, i.e., annotations exist more likely for large families. Larger families may be easier to predict than smaller ones. In fact, this may exactly explain why proteins from the largest families tend to be predicted better by pLM-than by MSA-based methods ^5,11^ which clearly contain more information ^45^. Thus, our proxy (Fig. 1: “no MSA”) may not capture the full story. Nevertheless, the fraction of proteins most interesting for the novelty-value they carry is likely to be substantially higher for understudied pathogens and species than for the human proteome. In other words, for many proteins that might become relevant for health and biotechnology, predictions not using MSAs may provide much better answers than the state-of-the-art based on evolutionary information.

### Do mutation movies capture any aspect of structural plasticity?

One important direction of protein research is the “hallucination of new structures”^46^. Could the mutation movies we introduced here help to trace evolution in the tube (e.g., the evolution from one shape to another through sequence variation), or to find new solutions fitting to particular constraints? More basically: do differences between mutant and wild type capture any aspects of the free energy landscape of proteins^47^? If so, would those aspects matter for understanding the problems of protein dynamics theoretically and/or would they have any practical impact? The correlation between functional DMS data and differences in structures predicted for mutants appeared encouraging, in particular, given that the DMS assays function, i.e., even experimentally measured structural differences may not result in higher levels of correlation.

*EMBER3D* captured effects of sequence variation upon structure better than methods such as *AlphaFold2* although it predicts structures for wild types with sufficient evolutionary information better. This clearly implied one advance of our new technology toward understanding protein function.

## Conclusions

We introduced a novel structure prediction system, *EMBER3D*, computing both 2D (distance maps) and 3D structure (backbone coordinates) from sequence alone in milliseconds for average-length proteins using consumer-grade hardware, based on embeddings from protein Language Models (pLMs), in particular on ProtT5. While the accuracy of EMBER3D does not rival state-of-the-art systems with focus on quality, such as AlphaFold2, RoseTTAFold, or even ESMFold, our method is more sensitive to small changes in amino-acid sequence and therefore useful as a predictor for structural effects of point mutations (single amino acid variants, SAVs) or combinations thereof.

This ability, paired with its speed, allows EMBER3D to render almost-real-time *protein mutation movies* (PMM) that visualize the predicted structural effect of each possible SAV in a protein. Effectively, this enables live-editing of sequences with near-instantaneous feedback of the impact on the predicted structure, e.g., in a webserver. Despite its lower accuracy, EMBER3D predictions suffice for highly sensitive structure search through *Foldseek*, i.e., successful root the identification of proteins of similar structure through prediction. Many proteins relevant for novel discoveries might be in small families for which EMBER3D appears to predict not only faster but also better than AlphaFold2. EMBER3D’s lightning-fast structure predictions open the door to a variety of new use-cases and possibly could lead to finding mutation paths that switch proteins between different structural conformations.

## Methods

### Dataset

We obtained 102,823 sequences from SidechainNet ^48^ as the base for training and validation; SidechainNet adds torsion angles and coordinates for all heavy atoms to ProteinNet12 ^49^. Due memory constraints, we removed proteins >430 residues, yielding 93,286 sequences for training. To avoid bias from overrepresented families, we clustered the training set with *UniqueProt* ^50^ at the default Hval=0. Following AlphaFold developers ^4^, we trained on randomly picked samples from the resulting 11,580 clusters. We optimized hyperparameters on the in-built validation set of SidechainNet with 39 CASP12 ^51^ proteins. We assessed the performance of the final model on 18 CASP14^52^ domains from free-modeling (FM) and template-based modeling hard (TBM-hard).

### ProtT5 protein language model

In this work, we generated embeddings for each protein sequence using the pLM ProtT5-XL-UniRef50 ^11^ (for simplicity ProtT5) built in analogy to the NLP model T5^53^. ProtT5 was exclusively trained on unlabelled protein sequences from BFD (Big Fantastic Database; 2.5 billion sequences including meta-genomic sequences) ^54^ and UniRef50 ^55^. Ultimately, this allowed ProtT5 to learn constraints imprinted by structure and function upon protein sequences. As ProtT5 used only unlabelled sequences and we added no supervised training or fine-tuning, there was no risk of information leakage or overfitting to the tasks addressed by EMBER.

### Model architecture

We adopted a similar approach as *RoseTTAFold* ^42^ by using 2-track blocks, processing 1D and 2D features, followed by a series of 3-track structure blocks, jointly processing 1D, 2D and 3D information (Fig. S1). In contrast to RoseTTAFold, we used only one 2-track and two 3-track blocks to minimize runtime and memory.

We also replaced the original SE(3) transformer from RoseTTAFold by a more efficient NVIDIA implementation ^56^. Reducing the number of feature channels in the 1D and 2D pipeline reduced memory further; this allowed processing longer proteins on GPUs with less vRAM. Besides those changes, the internal layout of the attention blocks in the 2- and 3-track blocks of EMBER3D was identical to RoseTTAFold.

### Input, output, training

While RoseTTAFold inputs MSAs and template information to the 1D and 2D parts of the network, we solely relied on features derived by ProtT5 from single protein sequences. We extracted 1D representations in the form of ProtT5 embeddings and 2D representations from the ProtT5 attention heads as described in more detail elsewhere ^5^.

EMBER3D predicts C-beta distograms (42 bins), anglegrams (omega & theta: 37 bins, phi: 19 bins), and 3D coordinates in PDB format for all backbone atoms (C, C-alpha, N, O). As others ^4,42^, we estimated confidence by predicting the lDDT score for each residue, and stored predicted lDDT scores in the b-value column of the output PDB file.

EMBER was trained with Adamax at a learning rate of 0.001 until the lDDT score of the 3D predictions on the CASP12 validation set showed no improvement over 10 epochs.

### Relaxing predicted structures

In addition to the backbone coordinates obtained directly from our network, we used pyRosetta ^57^ to relax 3D models based on the predicted C-beta distance maps and angle maps. We first generated 75 different decoys using short-, mid- and long-range distances from the predicted distograms at varying levels of distance probability thresholds (here: [0.05, 0.45]). We then computed quality estimates of lDDT scores using DeepAccNet ^45^ to select the best predicted fold.

### Performance measures and error estimates

We assessed performance of predicted 3D models using the template-modeling score (TM-score), computed by TM-align ^58^, and the local Distance Difference Test (lDDT) ^27^. Standard errors computed were computed as follows:

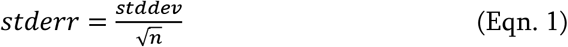

With *n* as the number of proteins, and *stddev* as the standard deviation obtained from NumPy ^59^. All error bars reflect the 95% confidence interval (CI95), i.e., 1.96 standard errors in results:

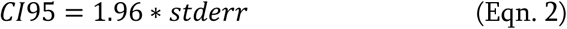

## Abbreviations used

2D: two-dimensional; 2D structure: inter-residue distances/contacts;
3D: three-dimensional;
3D structure: coordinates of atoms in a protein structure;
AH: attention heads;
BFD: big fantastic database;
CASP: Critical Assessment of protein Structure Prediction;
CNN: convolutional neural network;
DCA: direct coupling analysis;
DL: Deep Learning
DMS: deep mutational scanning;
ML: machine learning;
MSA: multiple sequence alignment;
pLM: protein Language Model;
PMM: protein mutation movie;
SAV: single amino-acid variant;
SCOP: Structural Classification of Proteins;
SOTA: state-of-the-art.

## Data availability

Pre-trained models, inference code, the mutation movie tool, a demo webserver and further resources are publicly available at https://github.com/kWeissenow/EMBER3D. A Google Colab notebook for structure prediction and mutation movie generation can be found at https://colab.research.google.com/drive/16qMVCRKPSLPI08vLxVZnBEB70qYKLqTV.

## Acknowledgements

We thank Tim Karl (TUM) for invaluable help with hard- and software and Inga Weise (TUM) for supporting many aspects of this work. Thanks to Minkyung Baek and the co-developers from Baker Lab for publishing the RoseTTAFold source code; thanks to Milo Mirdita (Seoul) for his contribution to making AlphaFold2 available through ColabFold. We gratefully acknowledge the support of NVIDIA with the donation of a Titan GPU used for development. Last not least, thanks to all who make their experimental data publicly available and all those who maintain such databases, in particular to Steve Burley and his team at the PDB.

## Notes

https://github.com/kWeissenow/EMBER3D

### Competing Interest Statement

The authors have declared no competing interest.

### Summary of Updates

Supplementary Materials have been updated. We added a link to a Google Colab notebook in the main text.

