## Supplementals for "Ultra-fast protein structure prediction to capture effects of sequence variation in mutation movies"

### Supporting online material for: Ultra-fast protein structure prediction to capture effects of sequence variation in mutation movies

**Konstantin Weissenow, Michael Heinzinger,**
**Martin Steinegger & Burkhard Rost**

#### Table of Contents for Supporting Online Material

#### Short description of Supporting Online Material

For brevity, we omitted the architecture of our model in the main text. Fig. S1 illustrates the conceptual layout of our approach.

We show that our proposed model reliably predicts its own performance (Fig. S2) and can be used as a rough pre-filter for more computationally expensive structure prediction systems.

We determined the influence of sequence length on prediction performance and show in Fig. S3 that accuracy drops only slightly for long proteins.

In Fig. S4 we show the convergence behaviour of our final model as well as an overfitting experiment on a single sample.

For the same overfitting experiment, we show a series of intermediate 3D predictions in Fig. S5.

Tables S1 and S2 provide detailed results for individual domains of our test set for all compared methods.

In Table S3 we compare model sizes of current structure prediction systems.

### Material

#### **Fig. S1:** Architecture overview

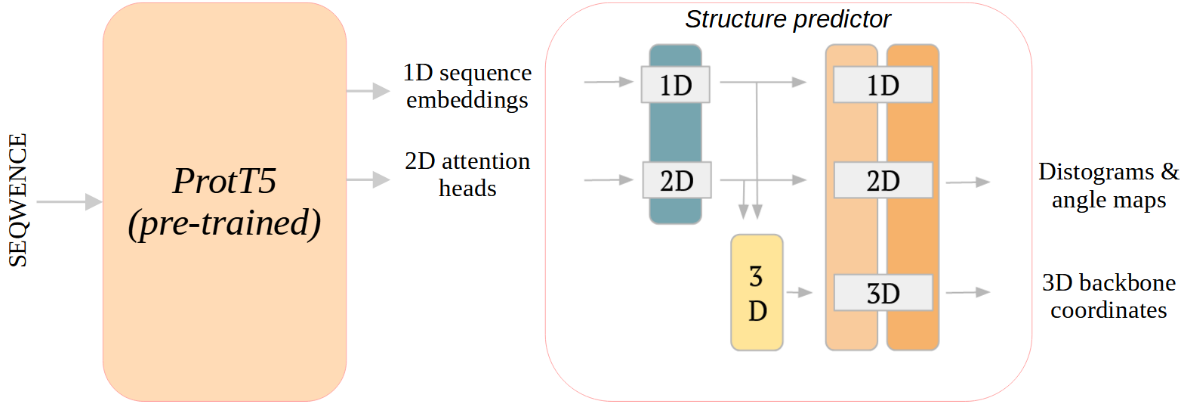

**Fig. S1: Architecture overview.** Our architecture consists of two major parts: our pre-trained protein language model ProtT5[1] (left; light orange box) is used to compute representations of the query sequence. We generate both a 1D representation from the last hidden layer of the model and extract 2D information from its attention heads (AHs). We feed both into the structure prediction network, which consists of a single 2-track block and two 3-track blocks nearly identical to the modules used by RoseTTAFold[2] (Methods). We predict 2D structure as C-beta inter-residue distance probability distributions (distograms) as well as angle-maps. The 3D part of the pipeline directly outputs backbone coordinates of the C, C-alpha, N and O atoms. We additionally predict lDDT[3] scores for each residue.

**Fig. S2:** EMBER3D reliably predicts its own performance

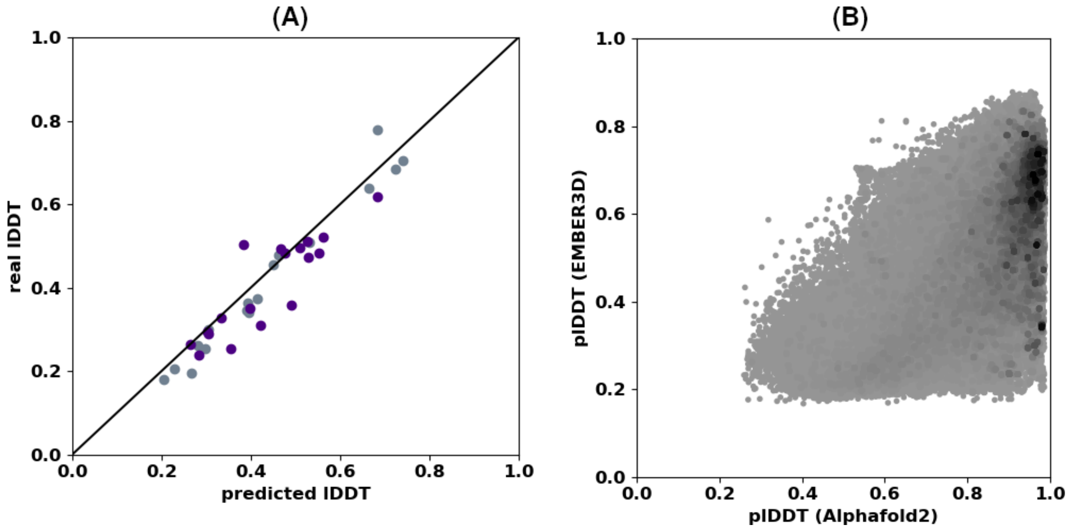

**Fig. S2: EMBER3D reliably predicts its own performance. (A)** EMBER3D predicted vs. experimental lDDT: x-axis: predicted lDDT[3] from EMBER3D; y-axis: lDDT computed against experimental structures. The closer to the diagonal (solid black line), the better the prediction. Dataset: all publicly released CASP14[4] targets, with the subset of samples representing our test set in violet. Visually the test set appeared similar to the entire set, although the largest outliers were observed in this subset. **(B)** On 538,488 proteins from Swiss-Prot[5], we correlated predicted plDDT confidence scores from AlphaFold2[6] (x-axis) to those from EMBER3D (y-axis). Coloring indicates point density (black: high density, and light grey: low). While EMBER3D is overall less confident, both methods still correlate with a Spearman rank of 0.53. Most high-quality AlphaFold2 predictions are also predicted with higher-than-average confidence by. Because of our method's high speed, it can be used as a very fast pre-filter on large datasets before running AlphaFold2 on proteins that are unlikely to result in a reliable prediction.

**Fig. S3:** Performance degrades only slightly for longer sequences

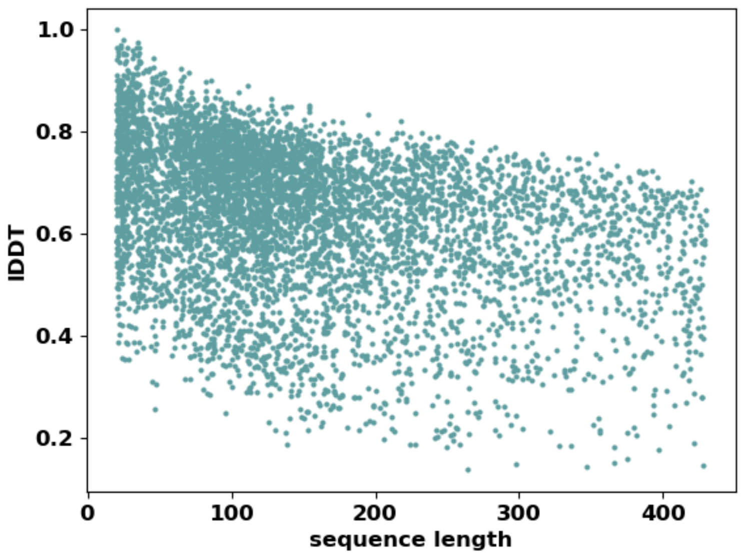

**Fig. S3: Performance degrades only slightly for longer sequences.** We measured the performance in lDDT[3] against experimental structures during the last epoch of training to evaluate the effect of protein length on the accuracy of our predictor on a large set of samples. Performance generally degrades for longer sequences, but not significantly (Spearman rank correlation: 0.37), which could be caused by overfitting on short samples. On our test set (CASP14 domains) we observed only a weak correlation (0.1), however most samples in the set are small (longest domain: 178 residues).

**Fig. S4:** Training dynamics: 2D prediction improves fast

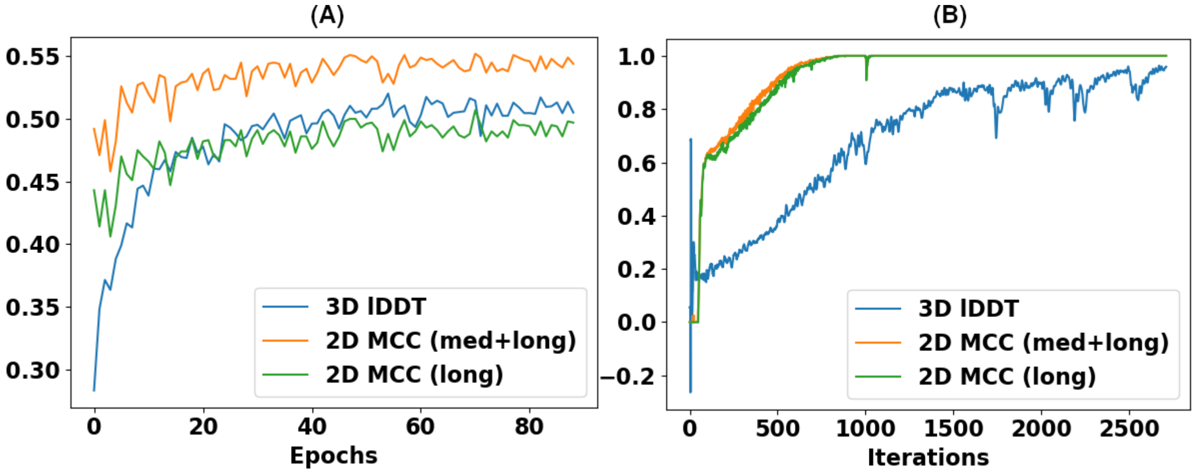

**Fig. S4: Training dynamics: 2D prediction improves fast.**
Training performance measured in C-alpha lDDT[3] on predicted backbone coordinates and Matthews Correlation Coefficient (MCC, Eqn. S1) on predicted contacts against experimental structures (**med+long**: medium and long-range contacts defined by a sequence separation of >4 residues; **long**: only long-range contacts; sequence separation of >23 residues):

$$MCC = \frac{TP*TN-FP*FN}{\sqrt{(TP+FP)(TP+FN)(TN+FP)(TN+FN)}} \quad \text{(Eqn. S1)}$$

For this, we obtained contact maps from predicted distograms by summing up probabilities of the distance bins representing distances below 8Å.
**(A)** Training dynamics of our final model on our validation set of CASP12[7] samples. As shown in our previous work, EMBER2[8], the AHs of our protein language model ProtT5[1] already suffice as a noisy contact predictor. Even after just one epoch of training, our model reaches reasonable prediction quality in its 2D output.
**(B)** Similarly, during an overfitting experiment on just a single sample (Ykud L,D-transpeptidase[9]; PDB 4a1k), our model is able to perfectly predict 2D structure within less than 1000 iterations, while reasoning over 3D geometry proves to be more challenging.

**Fig. S5:** Global structure is discovered before local structure

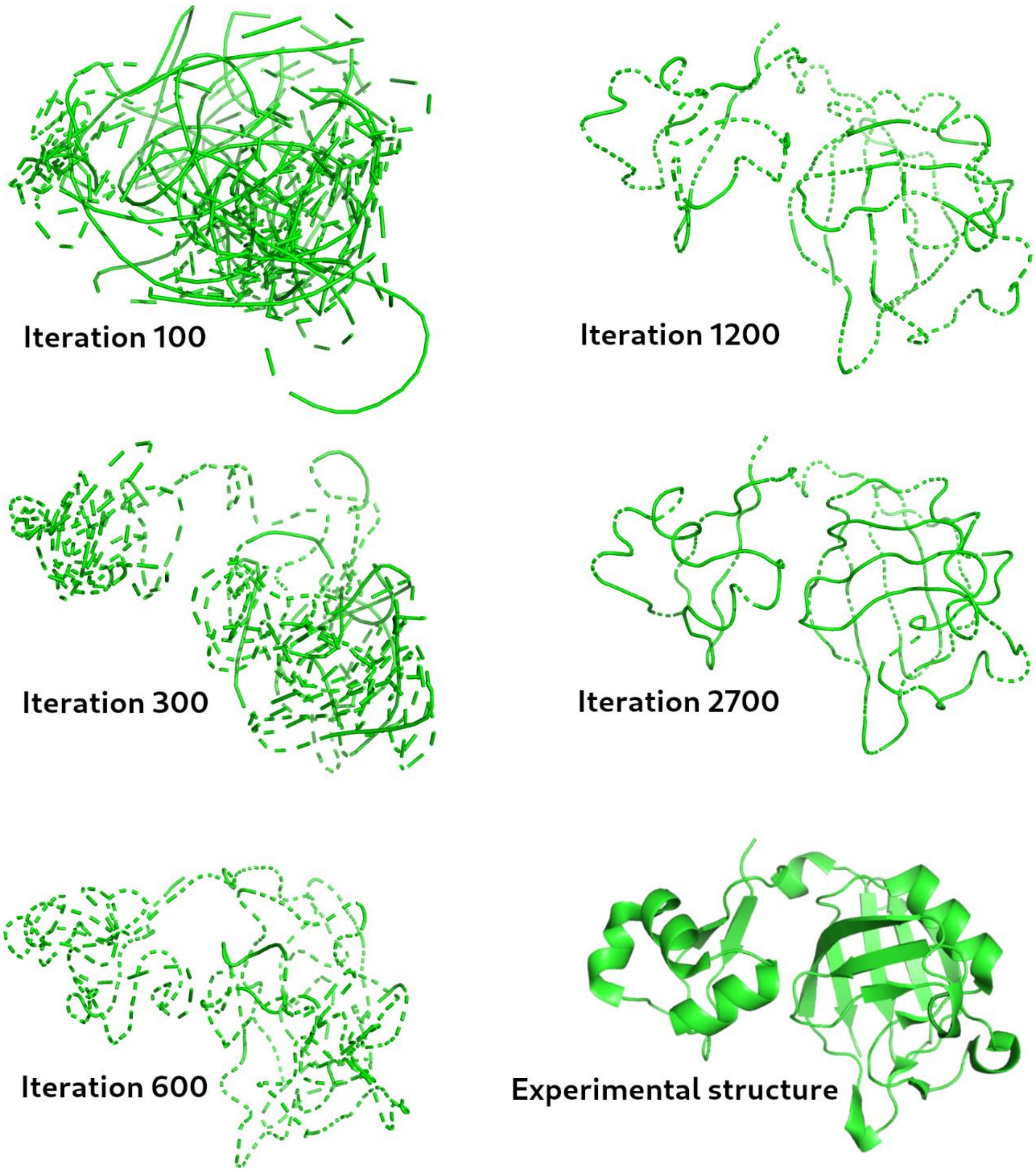

**Fig. S5: Global structure is discovered before local structure.** Intermediate 3D outputs of our model during the overfitting experiment evaluated in Fig. S4B (Ykud L,D-transpeptidase[9]; PDB 4a1k). After just 300 iterations, the predicted backbone geometry shows a clear separation of sub-units before optimizing more local structural conformations, e.g. secondary structure. We hypothesize this effect to be caused by the fast improvement of predicted inter-residue distances (MCC, Fig. S4B).

#### Table S1: TM-score on CASP14 test set. *

| | AF2 | AF2_single | OmegaFold | HelixFold | ESMFold | EMBER2 | EMBER3D | EMBER3D (UR) |
|---|---|---|---|---|---|---|---|---|
| Domain | | | | | | | | |
| **T1026-D1** | 0.77 | 0.56 | 0.97 | 0.85 | 0.95 | 0.74 | 0.74 | 0.71 |
| **T1027-D1** | 0.61 | 0.39 | 0.86 | 0.37 | 0.63 | 0.27 | 0.38 | 0.3 |
| **T1029-D1** | 0.53 | 0.51 | 0.46 | 0.42 | 0.5 | 0.33 | 0.32 | 0.33 |
| **T1030-D1** | 0.85 | 0.61 | 0.87 | 0.81 | 0.97 | 0.61 | 0.66 | 0.43 |
| **T1030-D2** | 0.51 | 0.78 | 0.92 | 0.55 | 0.87 | 0.58 | 0.7 | 0.42 |
| **T1031-D1** | 0.83 | 0.44 | 0.75 | 0.33 | 0.37 | 0.35 | 0.41 | 0.31 |
| **T1032-D1** | 0.7 | 0.24 | 0.78 | 0.63 | 0.7 | 0.62 | 0.71 | 0.62 |
| **T1033-D1** | 0.83 | 0.34 | 0.88 | 0.33 | 0.32 | 0.34 | 0.37 | 0.34 |
| **T1038-D1** | 0.93 | 0.3 | 0.34 | 0.33 | 0.36 | 0.27 | 0.32 | 0.34 |
| **T1038-D2** | 0.91 | 0.51 | 0.85 | 0.65 | 0.73 | 0.32 | 0.62 | 0.34 |
| **T1043-D1** | 0.23 | 0.31 | 0.31 | 0.33 | 0.26 | 0.3 | 0.33 | 0.3 |
| **T1046s1-D1** | 0.93 | 0.78 | 0.95 | 0.42 | 0.81 | 0.51 | 0.45 | 0.48 |
| **T1046s2-D1** | 0.94 | 0.47 | 0.94 | 0.51 | 0.94 | 0.77 | 0.8 | 0.32 |
| **T1049-D1** | 0.92 | 0.29 | 0.42 | 0.35 | 0.35 | 0.31 | 0.32 | 0.34 |
| **T1056-D1** | 0.95 | 0.31 | 0.82 | 0.77 | 0.86 | 0.67 | 0.65 | 0.28 |
| **T1064-D1** | 0.72 | 0.34 | 0.78 | 0.32 | 0.3 | 0.27 | 0.45 | 0.29 |
| **T1082-D1** | 0.97 | 0.59 | 0.95 | 0.53 | 0.91 | 0.61 | 0.57 | 0.5 |
| **T1099-D1** | 0.85 | 0.34 | 0.9 | 0.43 | 0.36 | 0.29 | 0.31 | 0.31 |

* Individual TM-scores for all evaluated methods. *EMBER3D (UR)* shows scores without pyRosetta[10] refinement.

#### Table S2: lDDT on CASP14 test set. *

| | AF2 | AF2_single | OmegaFold | HelixFold | ESMFold | EMBER2 | EMBER3D | EMBER3D (UR) |
|---|---|---|---|---|---|---|---|---|
| Domain | | | | | | | | |
| **T1026-D1** | 0.67 | 0.26 | 0.90 | 0.66 | 0.88 | 0.55 | 0.54 | 0.48 |
| **T1027-D1** | 0.67 | 0.51 | 0.81 | 0.30 | 0.62 | 0.37 | 0.39 | 0.29 |
| **T1029-D1** | 0.56 | 0.47 | 0.47 | 0.41 | 0.48 | 0.35 | 0.37 | 0.31 |
| **T1030-D1** | 0.94 | 0.75 | 0.94 | 0.93 | 0.97 | 0.74 | 0.86 | 0.50 |
| **T1030-D2** | 0.51 | 0.77 | 0.95 | 0.51 | 0.89 | 0.64 | 0.68 | 0.48 |
| **T1031-D1** | 0.80 | 0.39 | 0.69 | 0.34 | 0.39 | 0.40 | 0.42 | 0.35 |
| **T1032-D1** | 0.78 | 0.24 | 0.80 | 0.62 | 0.76 | 0.56 | 0.63 | 0.51 |
| **T1033-D1** | 0.82 | 0.42 | 0.84 | 0.34 | 0.45 | 0.45 | 0.44 | 0.36 |
| **T1038-D1** | 0.91 | 0.25 | 0.36 | 0.26 | 0.34 | 0.32 | 0.33 | 0.26 |
| **T1038-D2** | 0.89 | 0.45 | 0.83 | 0.48 | 0.66 | 0.62 | 0.62 | 0.50 |
| **T1043-D1** | 0.29 | 0.29 | 0.29 | 0.31 | 0.28 | 0.33 | 0.33 | 0.24 |
| **T1046s1-D1** | 0.96 | 0.77 | 0.97 | 0.49 | 0.79 | 0.55 | 0.55 | 0.47 |
| **T1046s2-D1** | 0.90 | 0.36 | 0.88 | 0.35 | 0.86 | 0.61 | 0.67 | 0.62 |
| **T1049-D1** | 0.88 | 0.27 | 0.37 | 0.28 | 0.32 | 0.28 | 0.30 | 0.29 |
| **T1056-D1** | 0.91 | 0.30 | 0.81 | 0.66 | 0.80 | 0.57 | 0.58 | 0.52 |
| **T1064-D1** | 0.58 | 0.31 | 0.75 | 0.25 | 0.29 | 0.30 | 0.29 | 0.26 |
| **T1082-D1** | 0.94 | 0.53 | 0.94 | 0.47 | 0.90 | 0.56 | 0.59 | 0.49 |
| **T1099-D1** | 0.84 | 0.45 | 0.89 | 0.46 | 0.46 | 0.39 | 0.46 | 0.33 |

* Individual lDDT scores for all evaluated methods. *EMBER3D (UR)* shows scores without pyRosetta[10] refinement.

**Table S3: Model sizes of structure prediction systems.** *

| Model | Parameters (trainable) | Parameters (total) |
|---|---|---|
| AlphaFold 2 | 93M | 93M |
| OmegaFold | 670M | 670M |
| HelixFold | 93M | 93M |
| ESMFold | 690M | 3.7B |
| EMBER3D | 4.7M | 1.5B |

* Overview of model sizes for various structure predictors. The total number of parameters can differ from the number of parameters trained on structure prediction because some approaches, i.e., ESMFold and EMBER3D, rely on pre-trained pLMs to encode protein sequences.
